# Metabolic-Pathway-Presence-Heatmap (MPPH): Constructing phylogenetic trees based on metabolic pathways

**DOI:** 10.1101/2023.06.27.546232

**Authors:** Yi-Heng Du, Jing-Hua Mu

## Abstract

Genome sequencing has revolutionized the study of biological systems, enabling exploration of species origins, evolution, and identification. However, traditional methods for constructing phylogenetic trees based on raw sequence data require substantial computational resources and may be challenging for biologists with limited computer knowledge. To address this, a lightweight tree-building tool was developed, Metabolic-Pathway-Presence-Heatmap (MPPH), leveraging Python programming and the KEGG metabolomics database to construct phylogenetic trees based on metabolic pathway information. This approach reduces computational and time requirements while focusing the analysis on metabolic pathways. The tool provides a rapid and reliable option for biologists to investigate the evolutionary and taxonomic aspects of species. Additionally, the tool incorporates a heatmap feature, allowing users to visualize the presence or absence of metabolic pathways across multiple species. The code is available at http://github.com/DeweyYihengDu/Metabolic-Pathway-Presence-Heatmap.

## Main

Over the last fifty years, sequencing technologies have revolutionized biological research, enabling the exploration of species origins, evolution, and identification (Heather and Chain, 2016). Heatmaps have become popular for visually representing data abundance in biology (Wu et al., 2018), providing a visual representation of data intensity and allowing researchers to identify patterns and correlations. They are widely used in genomics to analyze gene expression and other large datasets (Sun and Li, 2013; Gu, Eils and Schlesner, 2016). And phylogenetic trees are commonly used for species identification and evolutionary studies which depict the evolutionary relationships between organisms (Holmes, 2003). Various software programs, such as MEGA (Kumar et al., 2018), utilize sequence data to construct these trees using methods like neighbor-joining method (NJ) (Saitou and Nei, 1987), maximum-parsimony method (MP) (Mount, 2008), and maximum-likelihood method (ML) (Imbi Traat, 2013). And the phylogenetic trees are used in a wide range of applications including species identification. The integration of heatmaps and phylogenetic trees has emerged as a valuable approach in studying biological evolution. Heatmaps provide insights into data patterns, while phylogenetic trees offer a deeper understanding of species relationships (Allende, Sohn and Little, 2015). This combination can enhance our understanding of genetic and phenotypic evolution, uncovering the processes that shape the diversity of life.

Traditionally, the construction of phylogenetic trees, which depict the evolutionary relationships between species, has relied on the analysis of specific genetic markers, such as the 16S rRNA gene in bacteria (Clarridge, 2004; Watts et al., 2017). These markers provide a snapshot of the genetic diversity and relatedness between organisms. However, constructing phylogenetic trees based on these markers often necessitates dealing with large volumes of raw sequencing data, necessitating substantial computational resources and expertise. This poses a challenge for biologists who are not well-versed in computer science or lack access to high-performance computing infrastructure.

To address these challenges, a novel and lightweight tree-building tool has been developed, leveraging the power of Python programming and the comprehensive KEGG (Kyoto Encyclopedia of Genes and Genomes) metabolomics database (Kanehisa and Goto, 2000). This innovative approach capitalizes on the information about the presence or absence of metabolic pathways in species to construct phylogenetic trees. By focusing on metabolic pathways, the tool streamlines the computational requirements and significantly reduces the time and resources needed to generate accurate phylogenetic trees. In fact, the computation time is now reduced to a matter of minutes, enabling biologists to obtain rapid and reliable results without being impeded by computational limitations.

## Method

The tool eliminates the need for users to download raw data separately, as it seamlessly accesses the extensive resources of the KEGG database. Users can simply input the taxonomic information, such as the genus, family, order, or class name of the target species, and the tool will automatically retrieve the relevant data from the KEGG database (Figure 1). This streamlined process ensures that users can swiftly access the required information without being burdened by complicated data retrieval procedures.

**Figure 1.**
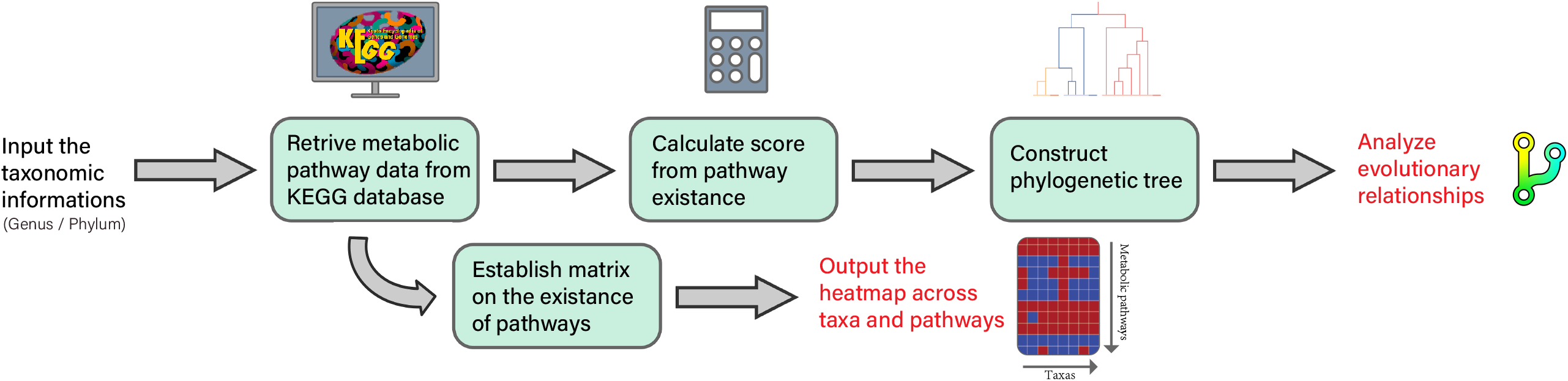
Workflow of Metabolic-Pathway-Presence-Heatmap (MPPH). The diagram shows the sequential steps involved in generating a metabolic pathway presence heat map (MPPH) and constructing a phylogenetic tree based on the presence or absence of pathways. The workflow starts by accessing the KEGG metabolomics database to retrieve species-specific pathway data. Next, the MPPH generates a heat map illustrating the presence (red) or absence (blue) of pathways across species. At the same time, the tool uses this pathway information to construct a phylogenetic tree representing the evolutionary relationships between species.

### Heatmap

The first essential feature of the Metabolic-Pathway-Presence-Heatmap (MPPH) tool is the heatmap, stands as a prominent highlight of the tool’s capabilities. It enables users to statistically analyze the presence or absence of metabolic pathways across one or multiple categories of multiple species (Figure 2). The resulting Heatmap visually represents the metabolic pathways, with red indicating their presence and blue indicating their absence. This intuitive representation allows users to readily observe and comprehend the distribution and patterns of metabolic pathways across species. Additionally, the tool generates output files in both PDF and PNG formats. The PDF files are editable, allowing users to annotate and tailor the data to their specific research requirements, thereby facilitating the identification of species and metabolic pathways of interest.

**Figure 2.**
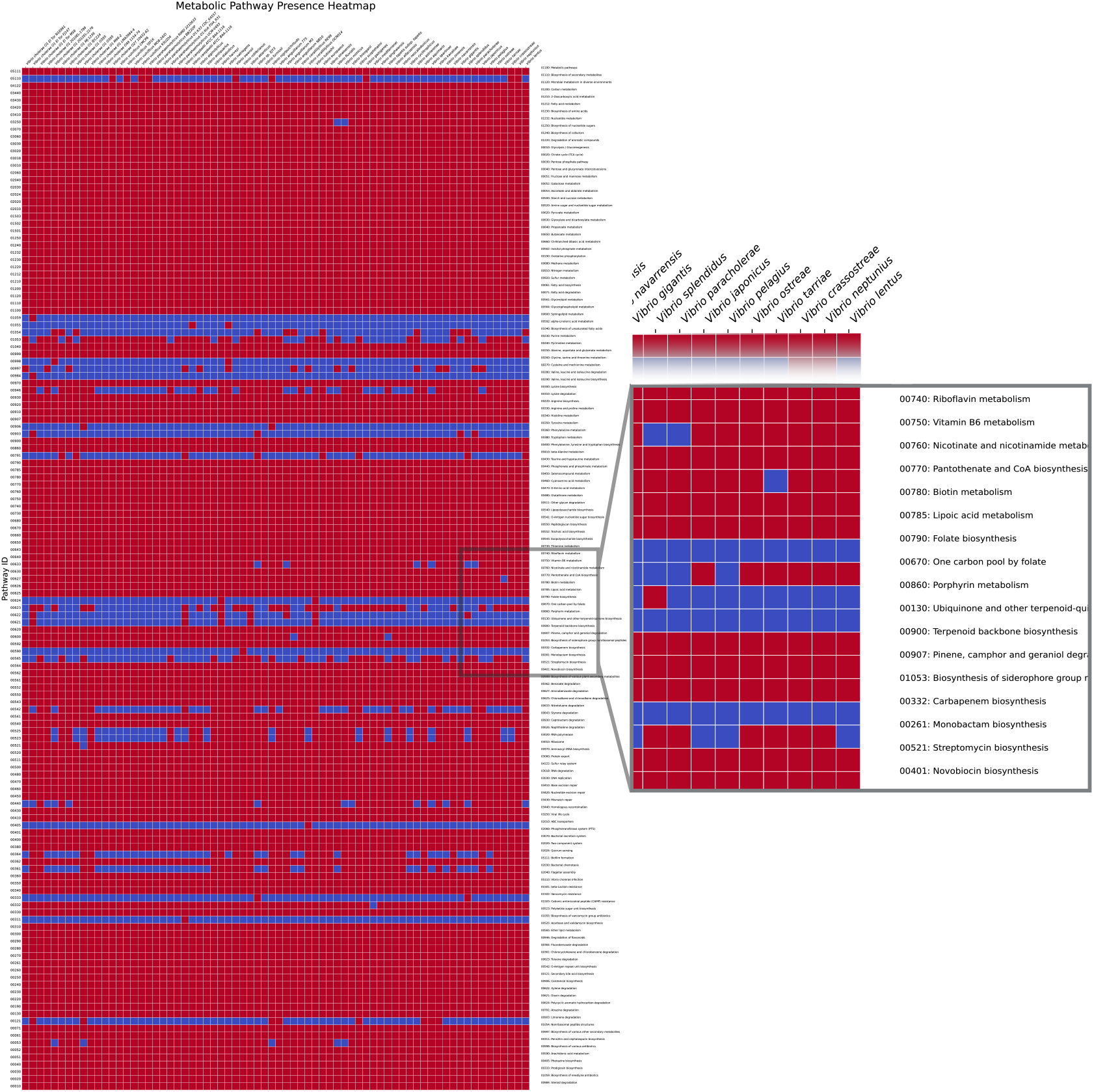
Heatmap plot of the presence or absence of metabolic pathways (example for *Vibrio* genus). The heat map represents the metabolic pathway profile of different *Vibrio* species, with red indicating the presence of a particular pathway and blue indicating its absence. The horizontal axis represents the species names of the bacteria, while the vertical axis represents the metabolic pathways of the bacteria.

Moreover, when generating the Heatmap, the tool simultaneously generates a CSV (Comma-Separated Values) file containing the underlying matrix data used for constructing the Heatmap. This matrix file includes species names and pathway identifiers, providing users with a comprehensive overview of the data. Importantly, users have the flexibility to manually modify the matrix, adding or removing entries as needed. This flexibility allows users to refine and customize the data for further analysis, while also promoting interoperability with other analysis tools and software platforms.

### Phylogenetic tree

The second feature of the tool is the phylogenetic tree, which serves as a visual representation of the evolutionary relationships between species based on the presence or absence of metabolic pathways. Building upon the matrix data generated earlier, the tool assigns scores to each species depending on the presence or absence of specific metabolic pathways. A score of 1 is assigned to indicate the presence of a pathway, while a score of 0 denotes its absence. These scores are then used to construct a phylogenetic tree. Species with similar pathway profiles cluster together, indicating a shared evolutionary history (Figure 3). The resulting tree provides a comprehensive overview of the relationships between species and offers insights into their evolutionary divergence and convergence.

**Figure 3:**
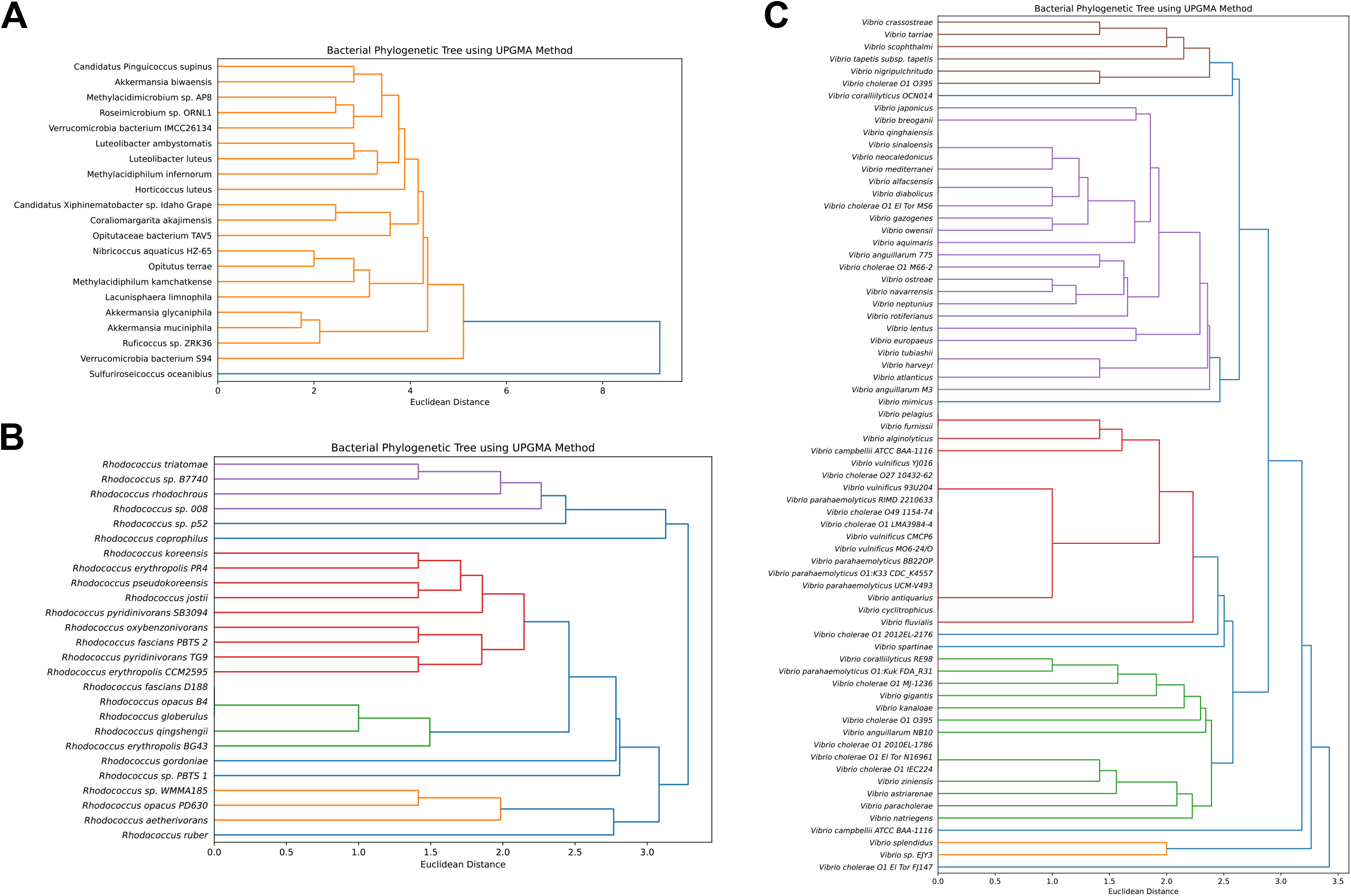
Phylogenetic Trees Based on Metabolic Pathways. **(A)** Phylogenetic tree of *Verrucomicrobia* bacteria constructed using the Metabolic-Pathway-Presence-Heatmap (MPPH) tool. **(B)** Phylogenetic tree of *Rhodococcus* genus constructed using the MPPH tool. **(C)** Phylogenetic tree of *Vibrio* genus constructed using the MPPH tool. The trees are based on the presence or absence of metabolic pathways and provide insights into the evolutionary relationships among the respective bacterial groups.

The development of this lightweight tree-building tool represents a significant advancement in the field of bioinformatics, specifically catering to biologists who may have limited expertise in computer science but seek efficient and reliable methods for exploring the evolutionary and taxonomic aspects of species. By leveraging Python programming and the wealth of data available in the KEGG metabolomics database, this tool empowers biologists to swiftly generate phylogenetic trees, centered on metabolic pathways, that offer valuable insights into the evolutionary relationships and functional capacities of different species.

## Discussion

The MPPH tool offers several significant advantages, primarily by saving substantial computational resources and focusing research on the dimension of metabolic pathways. By utilizing metabolic pathway information, the tool streamlines the analysis process and allows researchers to gain insights into the functional aspects of species. However, it is important to acknowledge that the tool currently has certain limitations that should be addressed in future developments.

Firstly, by emphasizing the analysis of metabolic pathways, the tool inherently overlooks valuable information at the gene and protein levels. Details such as gene mutations and protein modifications, which can be crucial for understanding the functional implications of metabolic pathways, may not be captured by the tool’s current approach (Fitz-James and Cavalli, 2022). Therefore, it is essential to consider integrating complementary methods or expanding the tool’s capabilities to encompass genetic and proteomic data. This would enable a more comprehensive understanding of the underlying molecular mechanisms and enhance the accuracy of the phylogenetic analysis.

Secondly, the construction of the phylogenetic tree solely based on the presence or absence of metabolic pathways oversimplifies the complexity of biological systems. Different metabolic pathways contribute to organismal function to varying degrees. For instance, pathways like the tricarboxylic acid cycle (TCA) (Wood, 1946) may have a greater impact on overall metabolism. In this regard, future optimization of the tool could involve developing a weighted matrix that accounts for the relative importance or abundance of specific metabolic pathways across all species. By considering pathway significance, the tool would generate more accurate phylogenetic trees that better reflect the evolutionary relationships among organisms.

Furthermore, expanding the tool’s capabilities to integrate additional biological data, such as functional annotations, pathway enrichment analysis, and comparative genomics, would enhance its overall utility. These enhancements would allow researchers to explore the interplay between genetic variations, protein functions, and metabolic pathways, providing a more comprehensive understanding of the evolutionary and functional aspects of species.

In conclusion, while the MPPH tool offers significant advantages by focusing on metabolic pathways and reducing computational requirements, there are areas for improvement. Future developments should aim to address the limitations, such as integrating genetic and proteomic data, accounting for pathway significance, and expanding the tool’s functionality. By addressing these concerns, researchers can benefit from a more holistic and nuanced analysis of the evolutionary and functional characteristics of species.

## Conclusion

In summary, the integration of Python programming, the KEGG database, and innovative computational approaches has culminated in the creation of a lightweight tree-building tool. This tool not only circumvents the need for extensive computational resources and expertise but also significantly reduces the time and effort required to construct phylogenetic trees. By capitalizing on metabolic pathway information, the tool provides a rapid and reliable option for biologists to investigate the evolutionary origins, relationships, and functional attributes of diverse species.

